# Replication-Deficient Whole-Virus Vaccines Against Cytomegalovirus Induce Protective Immunity in a Guinea Pig Congenital Infection Model

**DOI:** 10.1101/2025.02.04.636391

**Authors:** Mark R. Schleiss, Claudia Fernández-Alarcón, Craig J. Bierle, Adam P. Geballe, Alexey Badillo-Guzman, Christine E. Tanna, Kanokpan Tsriwong, Mark Blackstad, Jian Ben Wang, Michael A. McVoy

## Abstract

Vaccines are needed to prevent congenital cytomegalovirus (CMV) infections. This study used the guinea pig cytomegalovirus (GPCMV) model to examine replication-deficient whole-virus vaccines against maternal viremia and congenital infection. Two recombinant GPCMVs, GP51-DD and GP52-DD, were engineered with destabilization domains fused to essential viral late proteins GP51 and GP52. These viruses, predicted to replicate in the presence of the synthetic ligand Shield-1 but not in its absence, were evaluated for Shield-1-dependence *in vitro* and for safety, immunogenicity, and efficacy in a pregnancy/challenge model. GP52-DD was profoundly Shield-1-dependent, producing no detectable infectious progeny in its absence. In contrast, the replication of GP51-DD was delayed in the absence of Shield-1, but the virus reached similar peak titers with or without the compound. GPCMV-seronegative guinea pigs received two subcutaneous injections of placebo (sham-immunized), GP51-DD, GP52-DD, or wild-type GPCMV (WT-GPCMV). DNAemia post-vaccination was detected in only one of 14 dams in each of the GP51-DD- and GP52-DD-immunized groups compared to 10/10 control animals immunized with WT-GPCMV, confirming the recombinant viruses were attenuated. GPCMV-specific antibody responses were similar in all three vaccinated groups. When immunized dams were bred and challenged with virulent GPCMV, DNAemia was detected in all sham-immunized controls and in 44% of GP52-DD-immunized dams (at significantly reduced levels) but was absent in dams immunized with GP51-DD or WT-GPCMV. Notably, immunization with GP52-DD, GP51-DD, or WT-GPCMV significantly reduced congenital GPCMV transmission (protective efficacies of 89%, 94%, and 100%, respectively). Thus, the replication-deficient GP52-DD vaccine was attenuated and protected against intrauterine GPCMV transmission.

**Importance:** Congenital cytomegalovirus (CMV) infections could be prevented by a vaccine, but most of the vaccine designs that have advanced to clinical trials have been subunit vaccines. These viral envelope glycoprotein-targeting designs have not yielded an effective vaccine that provides durable immunity. A vaccine that confers immune responses to a broader repertoire of viral immunogens, such as vaccines based on the whole virus, could enhance protection. However, there are concerns about the safety of live attenuated CMV vaccines. Using the guinea pig/guinea pig cytomegalovirus model of congenital infection, this study demonstrates that two replication-deficient whole virus vaccines are attenuated in animals while also being highly immunogenic, providing protective immunity against maternal viremia and fetal infection at levels similar to that conferred by a wild-type virus.

## 1. Introduction

Human cytomegalovirus (HCMV) is a major cause of disability in newborns, and the development of an effective vaccine is a major public health priority (1–3). Because CMVs are highly species-specific, preclinical vaccine development must utilize animal viruses that are closely related to HCMV. While murine and rhesus macaque CMVs have been used for this purpose, guinea pig cytomegalovirus (GPCMV) is a particularly useful small animal model for vaccine development since the virus, unlike murine CMV, will infect the placenta and fetus after primary maternal infection during pregnancy (4–7). Thus, the guinea pig/GPCMV model has been utilized to examine a variety of vaccine designs and antiviral therapy strategies towards the goal of developing interventions that can prevent congenital HCMV infection (8–15).

HCMV vaccine development has largely focused on subunit vaccines based on glycoprotein B (gB), which mediates membrane fusion and entry into all cell types, and the pentameric complex (PC), a target for potent antibodies that selectively neutralize HCMV entry into endothelial and epithelial cells (3, 16–18). A recent vaccine candidate, V160, utilized a replication-defective recombinant virus where essential viral proteins were fused to the FK506-binding protein 12-destabilization domain (DD; (19–21)). V160 requires the presence of the synthetic ligand Shield-1, which does not exist in nature, to stabilize viral-encoded DD-fusion proteins and replicate in cell culture (21, 22). In V160, HCMV immediate early 1 (IE1), immediate early 2 (IE2), and pUL51 proteins are fused to DDs (21). However, the degradation of IE1 and IE2 proteins, which helps eliminate viral replication, likely also limits the production of viral antigens with early or late expression kinetics *in vivo* and thus may impair the immunogenicity of the vaccine.

To optimize a DD-based, replication-defective vaccine strategy in the GPCMV model of congenital infection, we generated two Shield-1-dependent recombinant GPCMVs, designated GP51-DD and GP52-DD, where the essential late viral proteins GP51 or GP52 were fused to DDs. We evaluated the Shield-1-dependence of these viruses *in vitro*, and tested their safety, immunogenicity, and efficacy against congenital GPCMV infection in a vaccination/pregnancy/challenge model. We found that the GP51-DD and GP52-DD viruses were differentially dependent on Shield-1 for replication in tissue culture. When used as replication-deficient vaccines, both viruses were highly attenuated and immunogenic in guinea pigs and protected against maternal and congenital GPCMV infection at levels comparable to prior infection by replication-competent GPCMV.

## 2. Materials and Methods

### 2.1 Virus and cells

Viruses were propagated in GPL guinea pig lung fibroblasts (ATCC CCL-158, also known as JH4) maintained in F-12 medium and supplemented with 10% fetal calf serum (FCS, Fisher Scientific), 10,000 IU/l penicillin, 10 mg/l streptomycin (Gibco-BRL), and 0.075% NaHCO_3_ (Gibco-BRL). Growth curves, viral titers, and neutralization assays were performed as described previously (23). GPCMV stocks were prepared and titered by plaque assay using GPL cells as previously described (24). Wild-type GPCMV (WT-GPCMV) consisted of a salivary gland-derived stock (serially passaged 24 times in strain 2 guinea pigs) correspond to GPCMV strain 22122, prepared as previously described (25). Shield-1 was the gift of Dai Wang (Merck Vaccines) and was used at a final concentration of 2 μM when cells were cultured for viral stock preparations. Titrations of GP51-DD and GP52-DD viruses were performed in the presence of 2 μM Shield-1.

### 2.2 Construction of GP51-DD and GP52-DD viruses

The recombinant viruses GP51-DD and GP52-DD were derived by modifying the infectious bacterial artificial chromosome (BAC) clone N13R10r129-TurboFP635 (26). This clone was modified from BAC N13R10, a complete genomic clone of tissue culture-adapted GPCMV strain 22122 (23). N13R10 was first modified to restore an intact *gp129* ORF, which encodes a subunit of the GPCMV PC, generating the BAC N13R10r129 (27), then further modified by insertion of a marker cassette encoding the red fluorescent protein (RFP) TurboFP635 to produce N13R10r129-TurboFP635 (26).

Two-step galactokinase-mediated recombineering was used as described previously (27, 28) to construct the viruses GP51-DD and GP52-DD. In step one, a *galK* cassette encoding galactokinase was inserted into BAC N13R10r129-TurboFP635 between the first and second codons of ORFs *GP51* or *GP52* by PCR amplification of plasmid pgalK using the oligonucleotide pairs listed under step 1 in Table 1. These PCR products were transformed into *E. coli* strain SW102 cells containing BAC N13R10r129-TurboFP635, followed by colony selection on Gal-positive plates and verification of clones containing the expected *galK* insertions by PCR and targeted sequencing, as described previously (29). In step two, oligonucleotide pairs (Table 1) were used to PCR amplify DD-encoding sequences from the plasmid pENTR4-FKBP12DD N-term A (AddGene, plasmid #17414), and the products were then transformed into SW102 cells containing the appropriate N13R10r129-TurboFP635 BACs with *galK* insertions in *GP51* or *GP52* that were constructed in step 1 (30). Colonies were then isolated using Gal counterselection plates and clones containing the desired DD-encoding insertions were verified by PCR screening and confirmed by targeted Sanger sequencing (28).

**Table 1.**
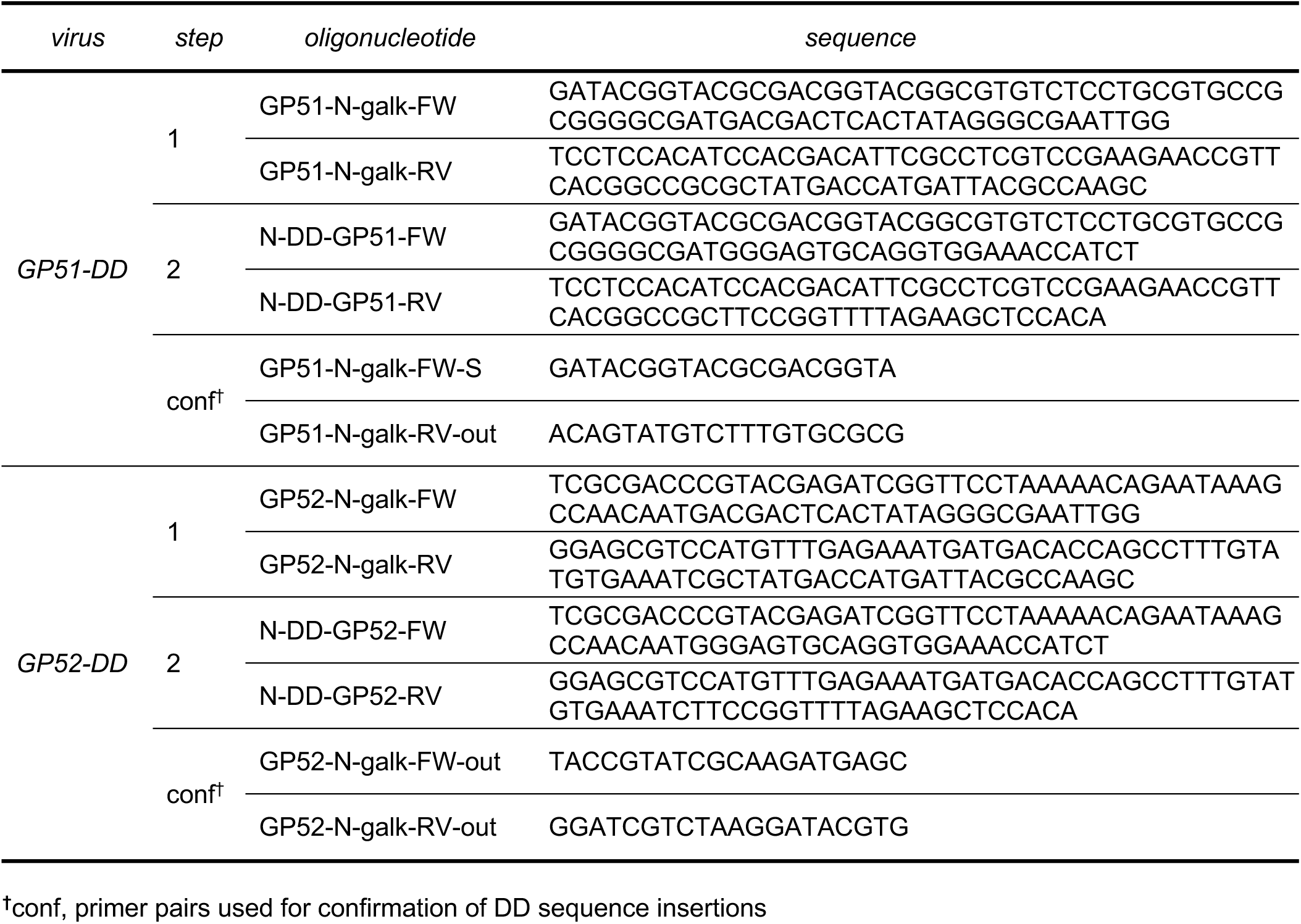
Oligonucleotides used for construction of GP51-DD and GP52-DD.

BAC-derived viruses GP51-DD and GP52-DD were reconstituted by transfection of BAC DNA into GPL cells using Effectene^®^ transfection reagent (Qiagen, Hilden, Germany), as previously described (31). Shield-1 (2 μM) was maintained in the culture medium during transfection and the subsequent generation of viral stocks. The correct genomic configuration of GP51-DD and GP52-DD viruses (Fig. 1A) was confirmed by PCR and sequence analyses of the DD domain sequences in the context of the *GP51* or *GP52* ORFs using the primer pairs listed as noted in Table 1.

**Fig. 1.**
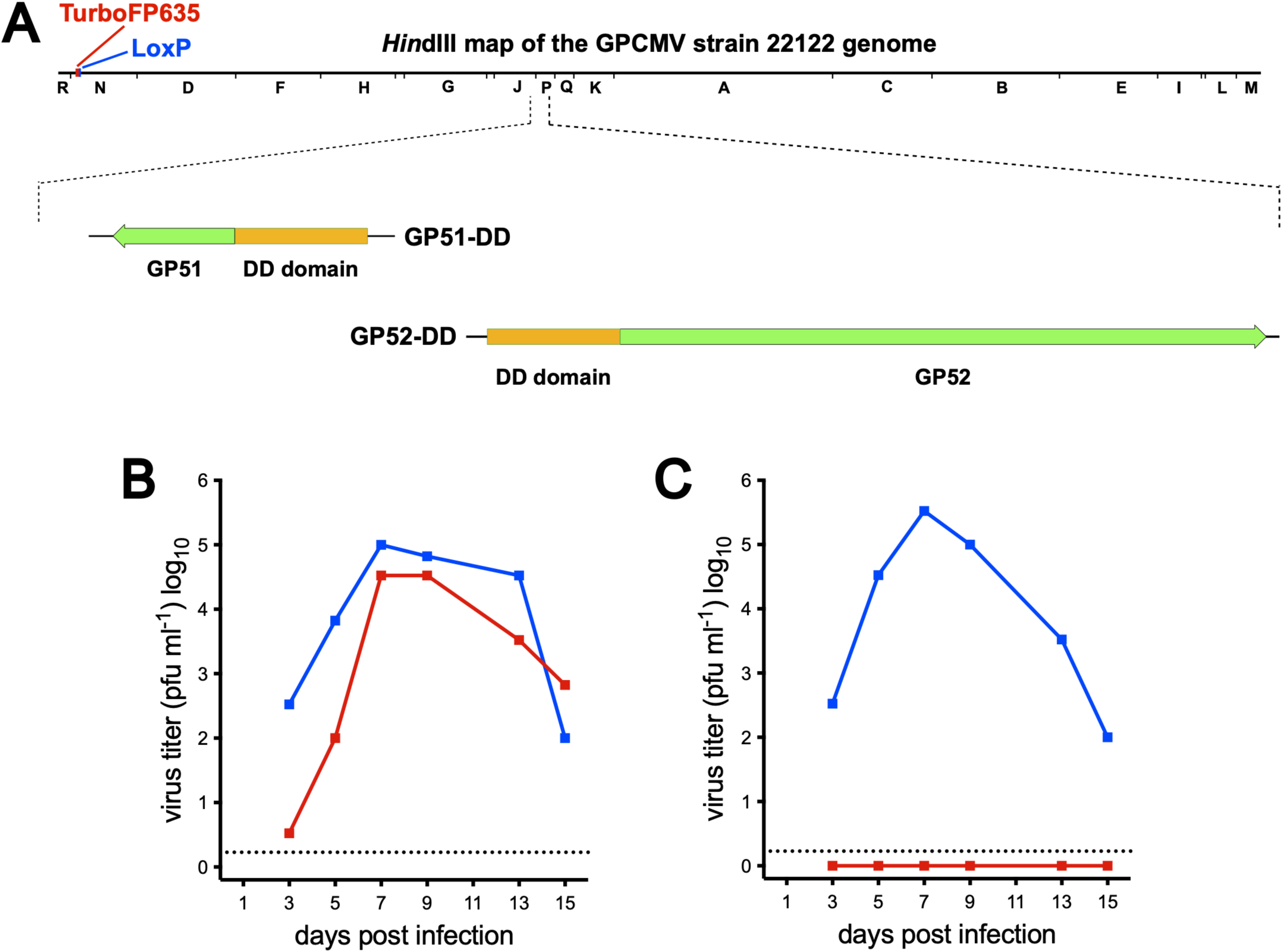
Design and *in vitro* growth properties of GP51-DD and GP52-DD recombinant GPCMVs. (A) *Hin*d III restriction map illustrating the locations of ORFs *GP51* and *GP52* in the GPCMV genome, as well as a marker gene cassette encoding the red fluorescent protein TurboFP635 and an adjacent *LoxP* site that is residual after cre excision of the BAC origin of replication. Expanded below are *GP51* and *GP52* ORFs that have been modified to encode N-terminal DD fusions to GP51 and GP52 in viruses GP51-DD and GP52-DD, respectively. Replicate GPL cultures were infected with (B) GP51-DD or (C) GP52-DD at an MOI of 0.01, then maintained for 15 days in medium either containing (blue) or lacking (red) 2 µM Shield-1. Samples of culture supernatants were collected on the days indicated and infectious titers were determined in the presence of 2 µM Shield-1. Dashed line indicates the LOD of 1.7 pfu mL^-1^.

### 2.3 In vitro viral replication assays

Growth curve analyses were performed as previously described (29). Replicate flasks of GPL cells were infected with GP51-DD or GP52-DD at an MOI of 0.01, and then maintained for 15 days in medium that either contained or lacked or 2 µM Shield-1. Samples of culture supernatants were collected at 3, 5, 7, 9 and 13 days post-inoculation and infectious titers were determined on fresh GPL cells in the presence of 2 µM Shield-1.

### 2.4 Ethics statement

All animal procedures were conducted in accordance with protocols approved by the Institutional Animal Care and Use Committee at the University of Minnesota (UMN; Protocol ID: 2206-40155A; approval date, September 13, 2022). Experimental protocols and endpoints were developed in strict accordance with the National Institutes of Health Office of Laboratory Animal Welfare (Animal Welfare Assurance # D16-00288), Public Health Service Policy on Humane Care and Use of Laboratory Animals, and United States Department of Agriculture Animal Welfare Act guidelines and regulations (USDA Registration # 41-R-0005). Guinea pigs were housed in a facility maintained by the UMN Research Animal Resources, accredited through the Association for Assessment and Accreditation of Laboratory Animal Care, International. All of the experimental procedures were conducted by trained personnel under the supervision of veterinary staff.

### 2.5 Guinea pig immunizations

Outbred Hartley guinea pigs (200-300 g) were purchased from Elm Hill Laboratories (Chelmsford, MA). Guinea pigs were confirmed to be GPCMV seronegative by enzyme-linked immunosorbent assay (ELISA), as previously described (32), prior to the initiation of the immunization studies. GPCMV-seronegative animals were randomly assigned to one of four vaccine groups. Guinea pigs were immunized subcutaneously (SC) twice at three-week intervals in the following group assignments: group 1 (N=14), GP51-DD vaccine (dose, 5 × 10^6^ PFU); group 2 (N=14), GP52-DD vaccine (dose, 5 × 10^6^ PFU); group 3 (N=10), WT-GPCMV, dose, 1.4 × 10^5^ PFU); and group 4 (N=10), sham-inoculated with PBS.

### 2.6 Immune assays

Blood samples were collected on days 7, 14, and 21 following each immunization; the day 21 bleed after the first dose of vaccine was obtained immediately before the second immunization was administered. Blood was also obtained from pregnant animals (described below, section 2.7) immediately prior to viral challenge; at days 7, 14, and 21 days post-challenge; and upon delivery.

GPCMV-specific IgG responses were reported as geometric mean titers (GMTs), determined as reciprocal of endpoint dilutions in an ELISA assay using GPCMV viral particles as coating antigen, as previously described (29). Samples negative at a 1:80 dilution were assigned a GMT value of 40 for statistical comparisons. GPCMV-reactive IgG avidity was measured as previously described (10). To determine the avidity of GPCMV-specific IgG responses, the ELISA described above was conducted using serum dilutions of 1:160, 1:320, 1:640, and 1:1280 in the presence or absence of 4 M urea. OD_450_ values for serum dilutions within the linear range of the ELISA assay (0.5–1.0 OD units) were used to calculate the avidity index for each sample, defined as the average of the ratios for different dilutions of the OD_450_ values obtained with 4 M urea to those obtained without urea.

A recombinant GPCMV expressing RFP, N13R10r129-RFP, was used to assay for neutralizing antibody responses (24). Neutralizing titers were reported as reciprocal GMTs of the lowest serum dilutions that resulted in ≥50% reduction in the number of RFP-positive foci on GPL cells detected 72 hours post-inoculation. Samples negative at a 1:40 dilution were assigned a GMT value of 20 for statistical comparisons.

### 2.7 Breeding and challenge during pregnancy

To assay for protection against GPCMV infection during pregnancy, most immunized dams were bred with GPCMV seronegative males; dams that were not bred were retained for future immunological studies. Pregnancy was confirmed by a progesterone ELISA, palpation, and ultrasonography (33). Pregnant dams were challenged at approximately 35 days gestation with 1 × 10^6^ pfu of WT-GPCMV administered by subcutaneous injection. Maternal blood samples were obtained at 7, 14, and 21 days post-challenge and again at delivery. Pregnancy outcomes were monitored and pup blood and tissues (lung, liver, and spleen) were harvested within 24 hours of birth, following necropsy as previously described (29).

### 2.8 Real-time qPCR Analysis

Viral loads were quantified by qPCR as described previously (15). Briefly, DNA was extracted from either 100 μl of citrated blood or from 0.05 g of homogenized tissues using the QIAamp 96 DNA QIAcube HT Kit (Qiagen). Amplification primers GP83TM_F1 (5’-CGTCCTCCTGTCGGTCAAAC-3’) and GP83TM_R1 (5’-CTCCGCCTTGAACACCTGAA-3’) were used at a final concentration of 0.4 µM, while the *GP83* hydrolysis probe (FAM-CGCCTGCATGACTCACGTCGA-BHQ1) was used at 0.1 µM. PCR was performed and the data analyzed as previously described (15). DNAemia was expressed as the total number of genome copies per mL of blood. Viral loads in tissue were expressed as genome copies per mg of tissue. PCR assays were considered positive if both replicates demonstrated amplifiable DNA, or if one of two replicates were positive in two independent experiments. The limit of detection (LOD) of the tissue assay was 2 genome copies/mg tissue, and 200 copies/mL for the blood PCR, and negative values were assigned these values for statistical comparisons, as previously described (15).

### 2.9 Statistical analyses

GraphPad Prism (version 10.0) was used for statistical analyses. Pup mortality and transmission rate data were compared using Fisher’s exact test with one-sided comparisons. Log_10_-transformed antibody responses (ELISA titers, neutralization titers, and IgG avidity indices) were compared across groups and at individual time points by one-way analysis of variance (ANOVA) and t-test, as previously described (34). DNAemia, blood and visceral organ viral loads, and pup birth weight data were compared by analysis of variance (ANOVA).

### 2.10 Data availability

The RFP-tagged GPCMV, N13R10r129-RFP, was sequenced during the course of these studies and a GenBank Accession number obtained, PQ384583.1. This data is available at URL: https://www.ncbi.nlm.nih.gov/nuccore/PQ384583.1

## 3. Results

### 3.1 Generation and in vitro characterization of Shield-1-dependent GPCMVs

The effectiveness with which different proteins are destabilized by fusion to DD varies widely, as does the ability of Shield-1 to stabilize different DD fusions (22). Hence, creating a virus that replicates efficiently in the presence of Shield-1 and poorly or not at all in its absence may require the construction and characterization of several viruses encoding DD fusions to different essential viral proteins (35). The preclinical development of V160 involved evaluating at least 11 HCMV recombinants encoding DD fusions to different viral proteins; among these transgenic viruses an absolute requirement for Shield-1 for viral replication was only noted when the DD was fused to UL51 or UL52 (21). This observation led us to target GP51 and GP52, the GPCMV homologs of UL51 and UL52, for destabilization. GP51-DD and GP52-DD were designed such that N-terminal DDs were fused to each respective protein (Fig. 1A). PCR and Sanger sequencing was then used to confirm the insertions of the DD sequences (data not shown). To assess for Shield-1-dependent replication of the GP51-DD and GP52-DD viruses *in vitro,* growth curves were performed in the presence or absence of Shield-1 (Fig. 1B). GP51-DD replicated in the absence of Shield-1, reaching similar peak titers in both conditions, but its replication kinetics were delayed two to three days without Shield-1. In contrast, replication of GP52-DD was profoundly dependent on Shield-1, reaching peak titers of ∼10^6^ pfu/mL in the presence of Shield-1 with no infectious progeny detected in its absence (Fig. 1B).

### 3.2 GP51-DD and GP52-DD are attenuated in guinea pigs

To evaluate the virulence and immunogenicity of GP51-DD and GP52-DD, outbred Hartley guinea pigs (>350 grams, 25-31 days of age) were randomly assigned to one of four groups.

Guinea pigs were injected subcutaneously (SC) with GP51-DD or GP52-DD (n=14; 5 × 10^6^ PFU/injection); WT-GPCMV (n=10; 1.4 × 10^5^ PFU/injection); or sham inoculated with PBS (n=10). These inoculations were repeated after 21 days. Blood and serum were collected every seven days post-inoculation so that viral loads and anti-GPCMV humoral responses could be quantified using qPCR, ELISA, and (at 21 days post-challenge) neutralizing assays.

Immunization with GP51-DD, GP52-DD, or WT-GPCMV had no significant effects on guinea pig weight relative to animals that received sham inoculations, suggesting that the infectious doses used did not cause systemic illness (data not shown). To determine whether GP51-DD or GP52-DD replicate *in vivo*, qPCR specific to GPCMV *GP83* was used to quantify DNAemia at days 7, 14, or 21 after each inoculation. Seven days after the first dose, DNAemia ranging from 3.4 × 10^4^ to 5.4 × 10^5^ genome copies/mL was detected in 10 of 10 animals inoculated with WT-GPCMV. By days 14 and 21 post-inoculation 7 of 10 and 6 of 10 animals remained DNAemic, respectively, in the WT-GPCMV challenged group (Fig. 2A). In contrast, three of 14 animals inoculated with GP51-DD or GP52-DD were DNAemic at day 7 post-inoculation in each of these respective groups, with significantly lower mean levels of DNAemia compared to the WT-GPCMV group (p<0.0001, Fig. 2A). A fourth animal in the GP51-DD group had low-level DNAemia at day 21 following the first inoculation. Following the second inoculation with these viruses, three of 14 animals in the GP51-DD, one of 14 in the GP52-DD, and six of 10 animals in the WT-GPCMV groups were DNAemic at the time of analysis for at least one of the time points (days 7, 14 and 21) measured, although none of the viral loads compared in the respective groups were statistically significantly different across groups by ANOVA (Fig. 2B). In total, six of 14 (43%) of the GP51-DD-inoculated animals and four of 14 (29%) of the GP52-DD-inoculated animals had demonstrable DNAemia, albeit at levels marginally above the limit of detection of the assay.

**Fig. 2.**
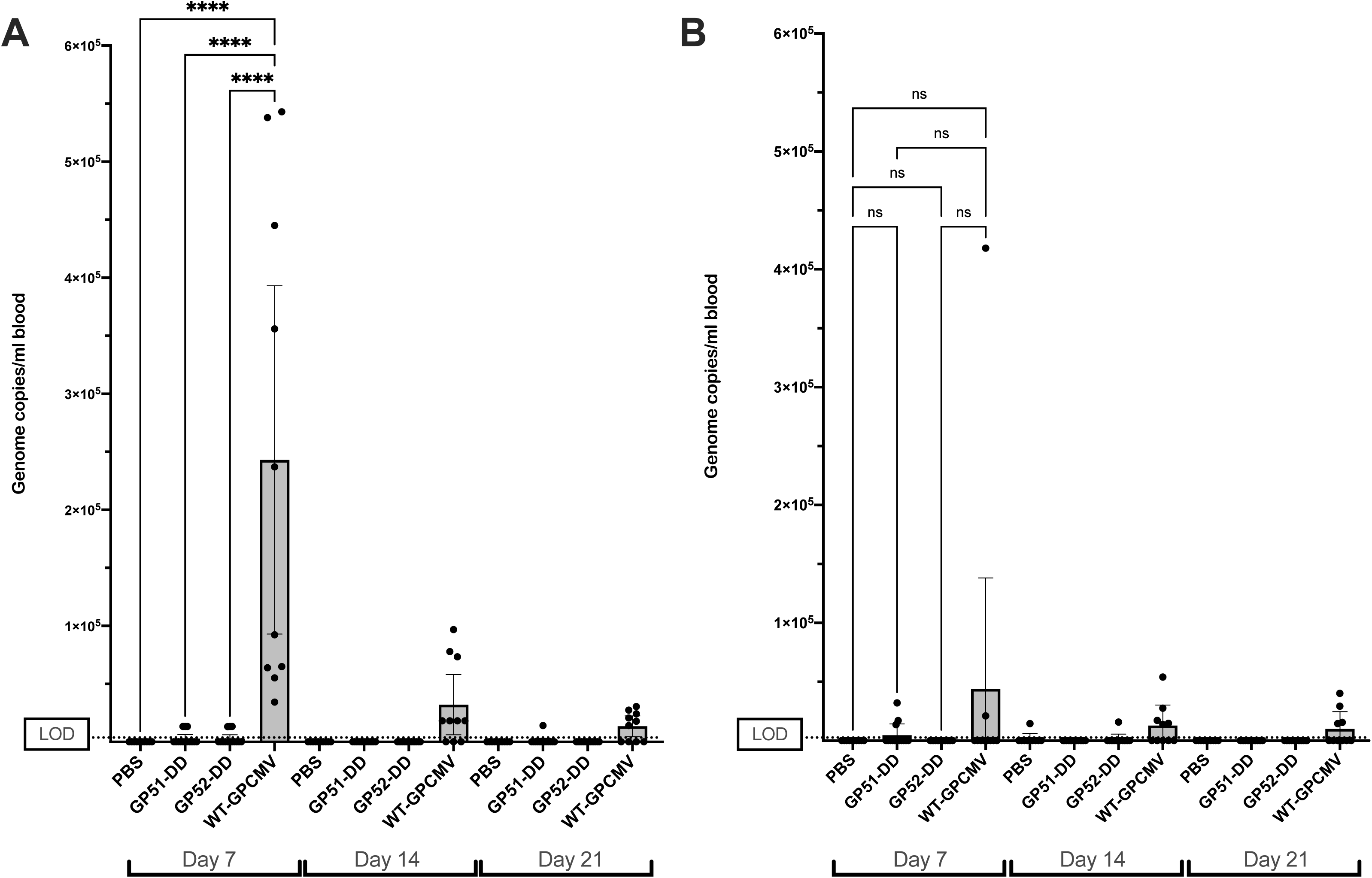
GP51-DD and GP52-DD are attenuated *in vivo* compared to wild-type GPCMV. Groups of female guinea pigs received two doses of GP51-DD or GP52-DD (n =14), replication-competent WT-GPCMV (n=10), or PBS (sham-inoculated) (n=10). Blood samples were collected 7, 14, or 21 days after the first (A) or second (B) inoculation and were assayed for GPCMV genome copy number by qPCR. Individual data points are shown with bars indicating means with their 95% confidence intervals (CI). Negative samples were assigned a value of 200 genome copies/ml of blood for statistical analyses, representing LOD of the assay (dashed line). By ANOVA, only the day 7 inoculation of wild-type virus resulted in a statistically significant difference in blood viral load, compared to the GP51-, GP52-, or PBS (sham) immunization (****p<0.0001).

### 3.3 GP51-DD and GP52-DD vaccines induce humoral responses comparable to those conferred by WT-GPCMV infection

Antibody responses were compared between animals that had been immunized with GP51-DD, GP52-DD, WT-GPCMV, or sham-inoculated with PBS. When GPCMV-reactive IgG titers were measured by ELISA, the GMTs were similar for animals immunized with GP51-DD, GP52-DD, and WT-GPCMV at seven and fourteen days after the first dose. At day 21, the IgG titers of WT-GPCMV-immunized animals were higher than those of animals immunized with either GP51-DD (p<0.0001) or GP52-DD (p<0.01, Fig. 3). This trend reversed following the second dose, where it was noted that the IgG titers at day 28 were significantly higher (p<0.0001) than those for animals immunized with GP51-DD or GP52-DD. The ELISA titer at day 35 in the vaccination sequence was also significantly higher in the GP52-DD vaccine group when compared to the WT-GPCMV group (p<0.05).

**Fig. 3.**
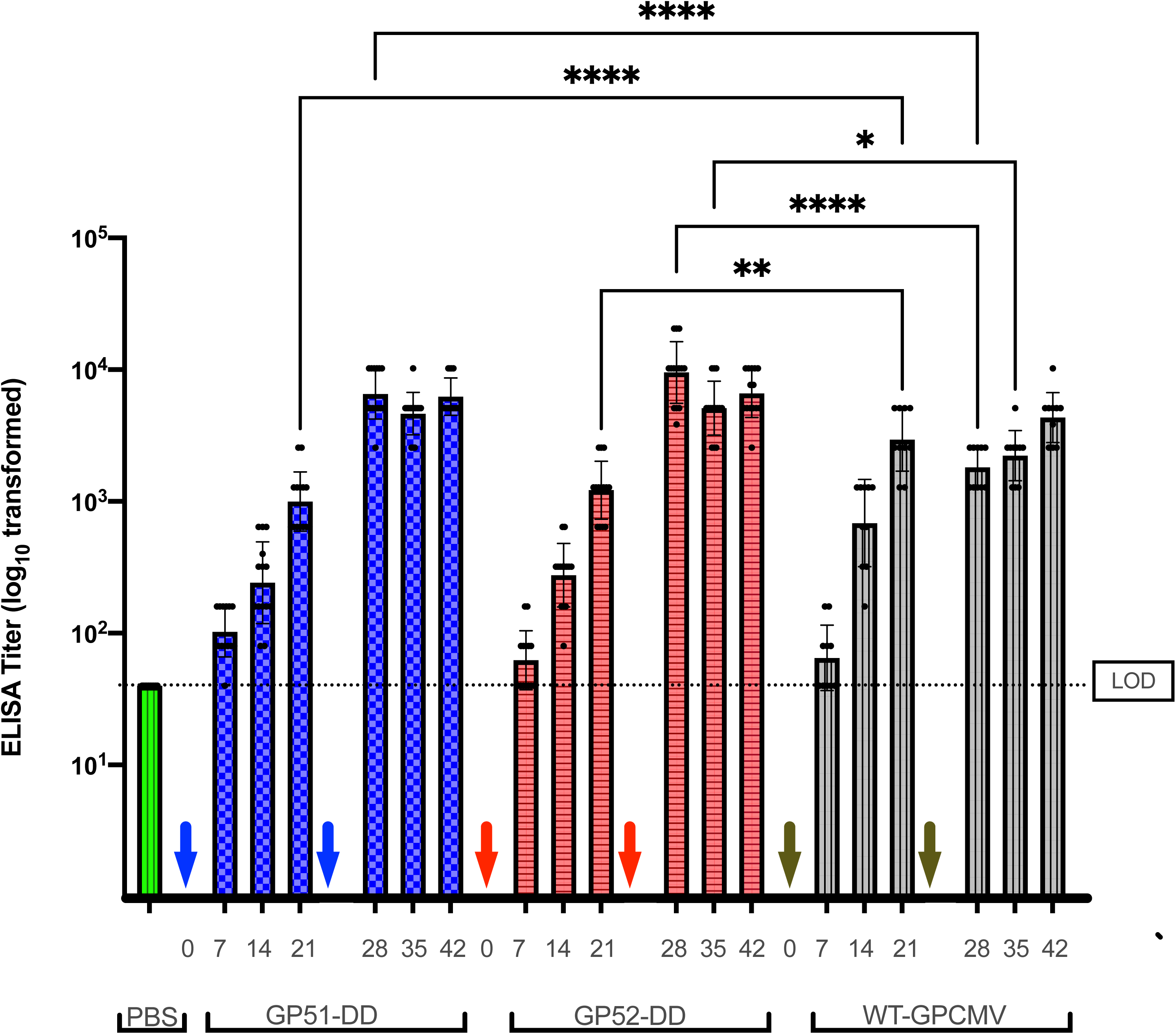
GP51-DD and GP52-DD vaccinations elicit similar IgG titers as WT-GPCMV vaccination. GPCMV-specific serum GMTs were measured by ELISA for GP51-DD (blue bars) and GP52-DD (red bars; n=14/group;), WT-GPCMV (gray bars; n=10), and PBS (sham-inoculated, green bar [n=10]) animals, day 42 time point only). All PBS immunized animals remained seronegative. Limit of detection for the ELISA is indicated by dashed line. Arrows indicate days of first and second inoculations; the day 21 bleed was just prior to administration of the second inoculation.

Inoculated guinea pigs were bred, and pregnancy was established in ten of ten dams immunized with GP51-DD, nine of ten dams immunized with GP52-DD, eight of eight dams immunized with WT-GPCMV, and ten of ten sham-inoculated controls (Table 2). Pregnant dams were challenged with WT-GPCMV at mid-gestation (approximately 35 days gestation) and blood was collected immediately before challenge, at 7, 14, and 21 days post-challenge, and after delivery. Since dams became pregnant asynchronously, viral challenge occurred variably between 65 and 98 days after the second vaccine.

**Table 2:** Group sizes^†^ and GPCMV-reactive IgG ELISA titers^‡^.

|  |  | **GP51-DD** |  | **GP52-DD** |  | **WT-GPCMV** |  | **PBS** |  |
| --- | --- | --- | --- | --- | --- | --- | --- | --- | --- |
|  | *day* | *N* | *titer* | *N* | *titer* | *N* | *titer* | *N* | *titer* |
| *1<sup>st</sup> dose* | 7 |  | 102 ± 45 |  | 59 ± 43 |  | 65 ± 48 |  | 40 ± 0 |
|  | 14 | 14 | 242 ± 206 | 14 | 276 ± 161 | 10 | 686 ± 477 | 10 | 40 ± 0 |
|  | 21 |  | 999 ± 673 |  | 1280 ± 771 |  | 3152 ± 1685 |  | 40 ± 0 |
| *2<sup>nd</sup> dose* | 28 |  | 6558 ± 2872 |  | 9547 ± 5741 |  | 1810 ± 675 |  | 40 ± 0 |
|  | 35 | 14 | 4637 ± 1869 | 14 | 5380 ± 2546 | 10 | 2229 ± 1121 | 10 | 40 ± 0 |
|  | 42 |  | 6241 ± 2400 |  | 6751 ± 2815 |  | 4331 ± 2262 |  | 40 ± 0 |
| *pre-challenge* | 0 | 10 | 2941 ± 1383 | 9 | 2765 ± 1582 | 8 | 4305 ± 2600 | 10 | 43 ± 13 |

<sup>†</sup>N, total animals per group
<sup>‡</sup>GMT ± SD

The IgG avidity index (AI) of GPCMV-reactive IgG in animals immunized with GP52-DD, GP51-DD, or WT-GPCMV was evaluated at the day 21 and 42 time points, and at delivery. AI increased progressively over the course of the experiment and no statistically significant differences were observed between vaccine groups (Fig. 4A). Sham (PBS)-immunized dams seroconverted to GPCMV challenge during pregnancy, and had measurable AIs by the time of delivery; mean AIs for the GP52-DD, GP51-DD, and WT-GPCMV groups were statistically significantly higher (p<0.0001) than that of the PBS group. Similarly, neutralizing antibody titers against GPCMV were not significantly different among groups immunized with two doses of GP51-DD, GP52-DD, or WT-GPCMV (Fig. 4B). Sham-immunized animals also developed neutralizing antibody following challenge with WT-GPCMV during pregnancy; however, post-delivery neutralizing potency at delivery was significantly lower than that observed for challenged dams that had undergone preconception vaccination with GP52-DD or WT-GPCMV (p<0.01). At day 42 post-preconception immunization, neutralizing titer in the WT-GPCMV inoculation group was also higher than that observed at delivery (post-challenge) in sham-immunized dams (p<0.05; Fig. 4B). Taken together, these findings suggest that vaccination with either GP51-DD or GP52-DD elicited humoral responses that were comparable to WT-GPCMV infection in magnitude of the reactive-IgG response, IgG AI, and neutralizing potency, despite the apparent restricted capacity of GP51-DD and GP52-DD to replicate *in vivo*.

**Fig. 4.**
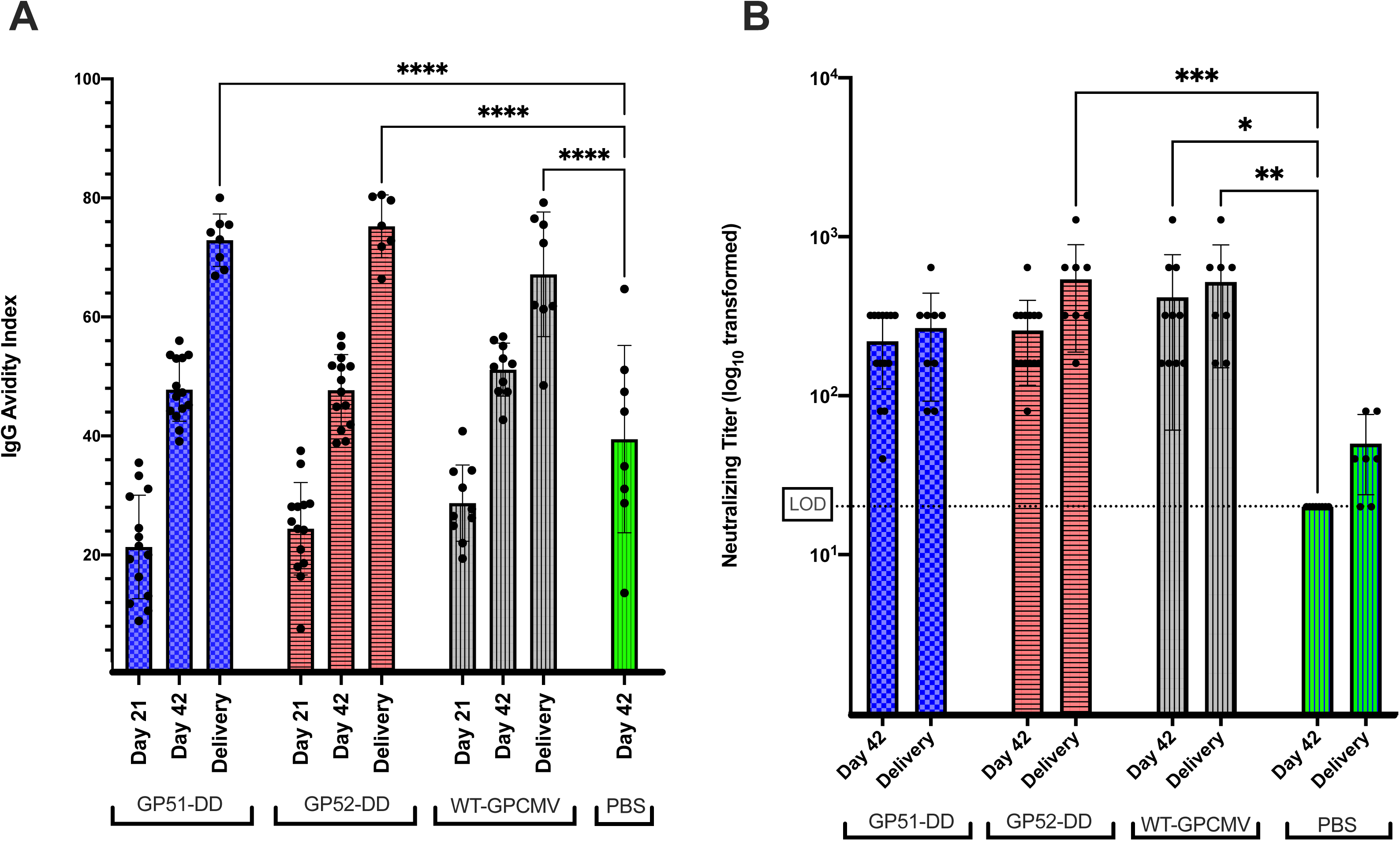
GP51-DD and GP52-DD vaccinations elicit similar IgG avidity and GPCMV neutralizing titers as WT-GPCMV vaccination. Guinea pigs were immunized as described previously and serum was collected 21 days after the first and second (day 42) doses, or after delivery following challenge during pregnancy. (A) Avidity indices for sera from animals inoculated with GP51-DD, GP52-DD, WT-GPCMV, or PBS (sham-inoculated) were determined (**** p<0.0001). (B) Neutralizing titers were determined after the second doses, or at delivery following challenge. Individual data points are shown with bars indicating GMTs ± one SD and levels of significance (*p<0.05; ** p<0.01; *** p<0.001). PBS samples refer to the control (sham-immunized) animals; only sera obtained post-pregnancy challenge, at the time of delivery, demonstrated a response (limit of detection of neutralization assay is indicated by dashed line).

### 3.4 Immunization with GP51-DD and GP52-DD protect against maternal viremia and congenital GPCMV infection

We next compared the protective efficacy of GP51-DD and GP52-DD relative to either sham-inoculation or immunization with WT-GPCMV. All sham-inoculated (PBS-vaccinated) dams were DNAemic at most time points post-challenge, while GPCMV DNA was not detected at any time point in any of the dams that were immunized with GP51-DD or WT-GPCMV. Of the nine GP52-DD-immunized dams, three were DNAemic on day seven post-challenge and low but detectable levels of DNAemia were detected in two of these animals at day 14. DNAemia was also detected in one other GP52-DD-immunized at delivery (Fig. 5, Table 3). Thus, four of 9 (44%) GP52-DD-immunized dams had DNAemia at one or more time points between WT-GPCMV pregnancy challenge and delivery. The overall impact of preconception vaccination has a highly significant impact on maternal viral load following pregnancy challenge. By ANOVA comparing all groups, the magnitude of DNAemia on day seven was significantly lower in the GP51-DD, GP52-DD, and WT-GPCMV-immunized dams (p<0.0001) compared to the sham-inoculated animals (Fig. 5).

**Fig. 5.**
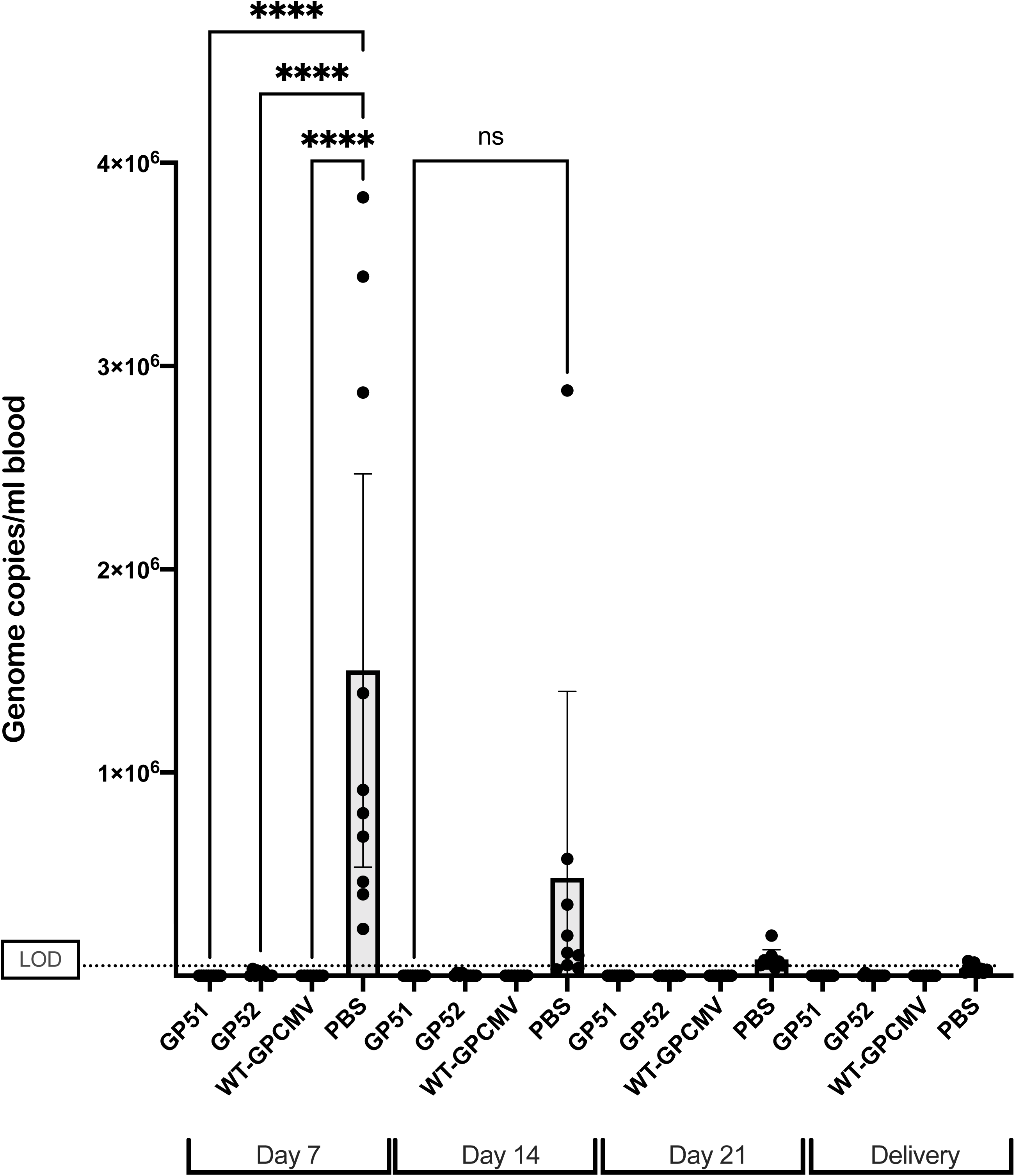
Vaccination reduces DNAemia in dams after GPCMV challenge during pregnancy. DNA was extracted from blood samples obtained from dams on days 7, 14, and 21 post-challenge, and at delivery. DNAemia was assayed by qPCR. Individual data points are shown with bars indicating means of log_10_ transformed data ± 95% CI (**** p<0.0001). Dashed line indicates limit of detection of assay.

**Table 3.**
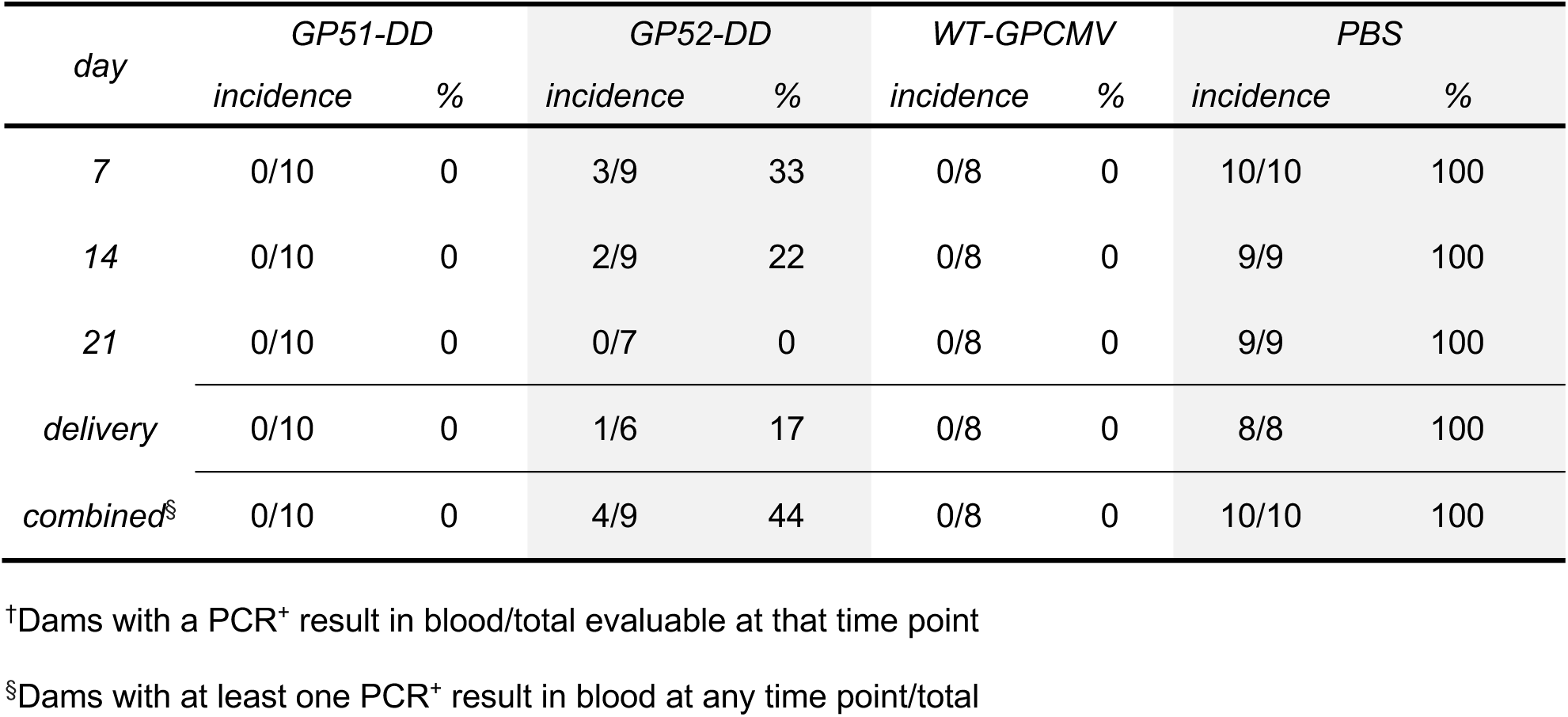
Incidence of maternal DNAemia^†^ following challenge of pregnant dams.

Next, the duration of pregnancies post-challenge with virulent WT-GPCMV challenge in dams, and the overall rates of pup mortality in the various groups, were compared (Table 4). No statistically significant differences were noted in pregnancy durations or pup mortality. However, the mean birth weights were lower for pups born to sham-inoculated dams, compared both to pups born to the WT-GPCMV group and pups born to GP52-DD-immunized group (Fig. 6; p<0.05).

**Fig. 6.**
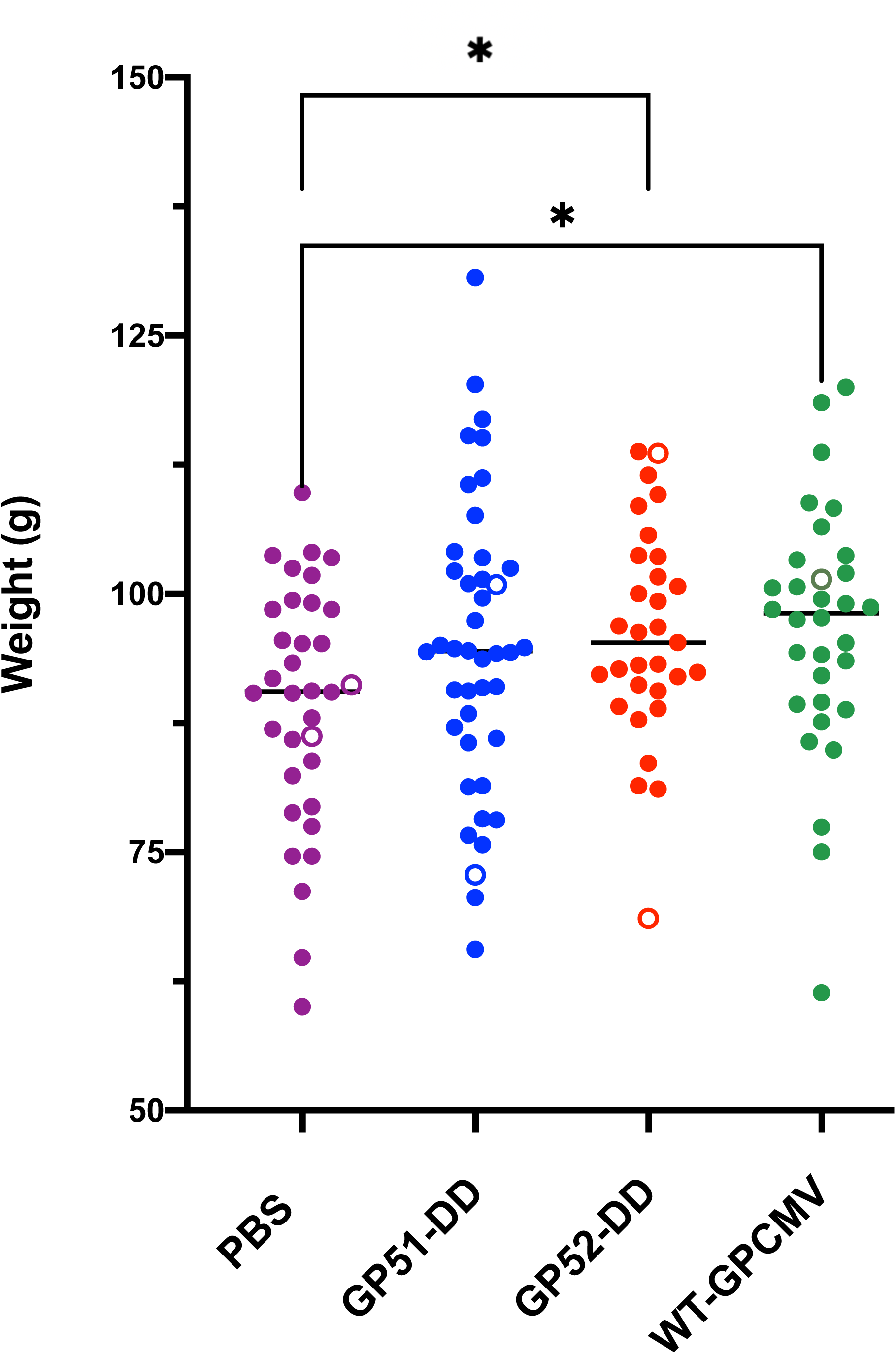
Pre-conception maternal vaccination results in improved pup birth weights after GPCMV challenge during pregnancy. Individual pup weights at birth are shown with maternal vaccine groups color-coded and stillborn pups indicated by open circles. For statistical comparison pups from dams that received GP51-DD or GP52-DD vaccines or WT-GPMV were analyzed in paired comparisons to the PBS (sham-inoculated) group by t-test with significant differences indicated (*p<0.05).

**Table 4.**
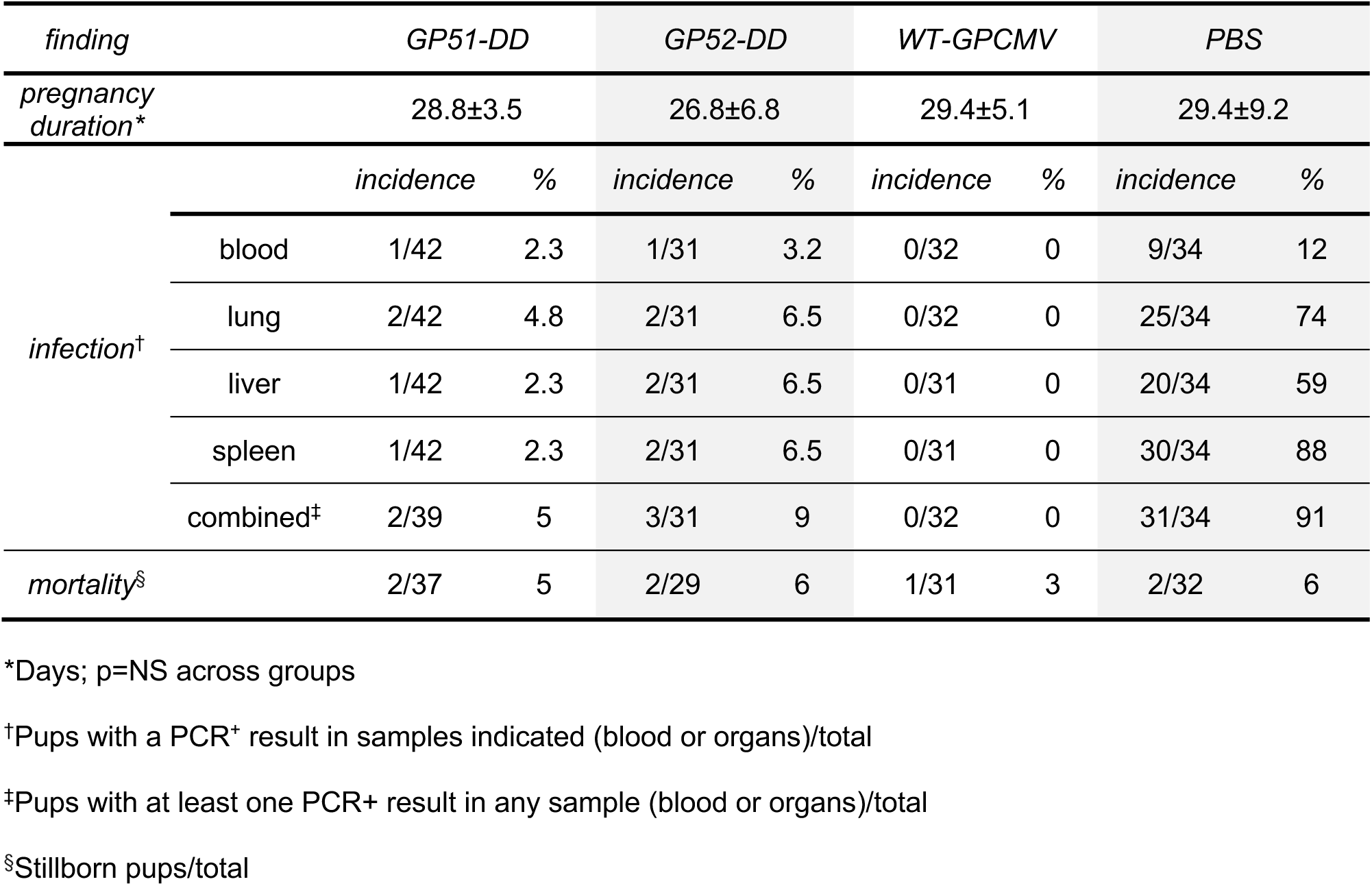
Pregnancy duration post-challenge, pup infection, and pup mortality across groups.

Finally, DNA was extracted from pup blood and tissues to assay for congenital infection (Table 4). 31 of 34 pups (91%) from sham-inoculated dams were infected, as evidenced by PCR detection of viral DNA in blood, lung, liver, or spleen. In contrast, viral DNA was observed in two of 39, three of 31, and none of 32 pups born to dams immunized with GP51-DD, GP52-DD, or WT-GPCMV, resulting in protective efficacies of 94%, 89%, and 100%, respectively (Fig. 7A, Table 4). Notably, one of the four DNAemic, GP52-DD-immunized dams gave birth to three pups, two of which had congenital GPCMV infection. Congenital infection did not occur in the three remaining DNAemic dams. A third pup in the GP52-DD group had congenital GPCMV infection (detected in blood only) although the corresponding maternal PCRs were all negative. Differences for each vaccine group by organ are shown in Fig. 7B. When viral loads were compared individually for lung, liver, and spleen, highly significant viral load reductions (p<0.00001) were noted for each vaccine approach compared to the sham (PBS)-immunized group. When viral loads were combined for all tissues in aggregate and then compared (Fig. 7B), significant overall reduction (p<0.0001) in pup viral load was also noted.

**Fig. 7.**
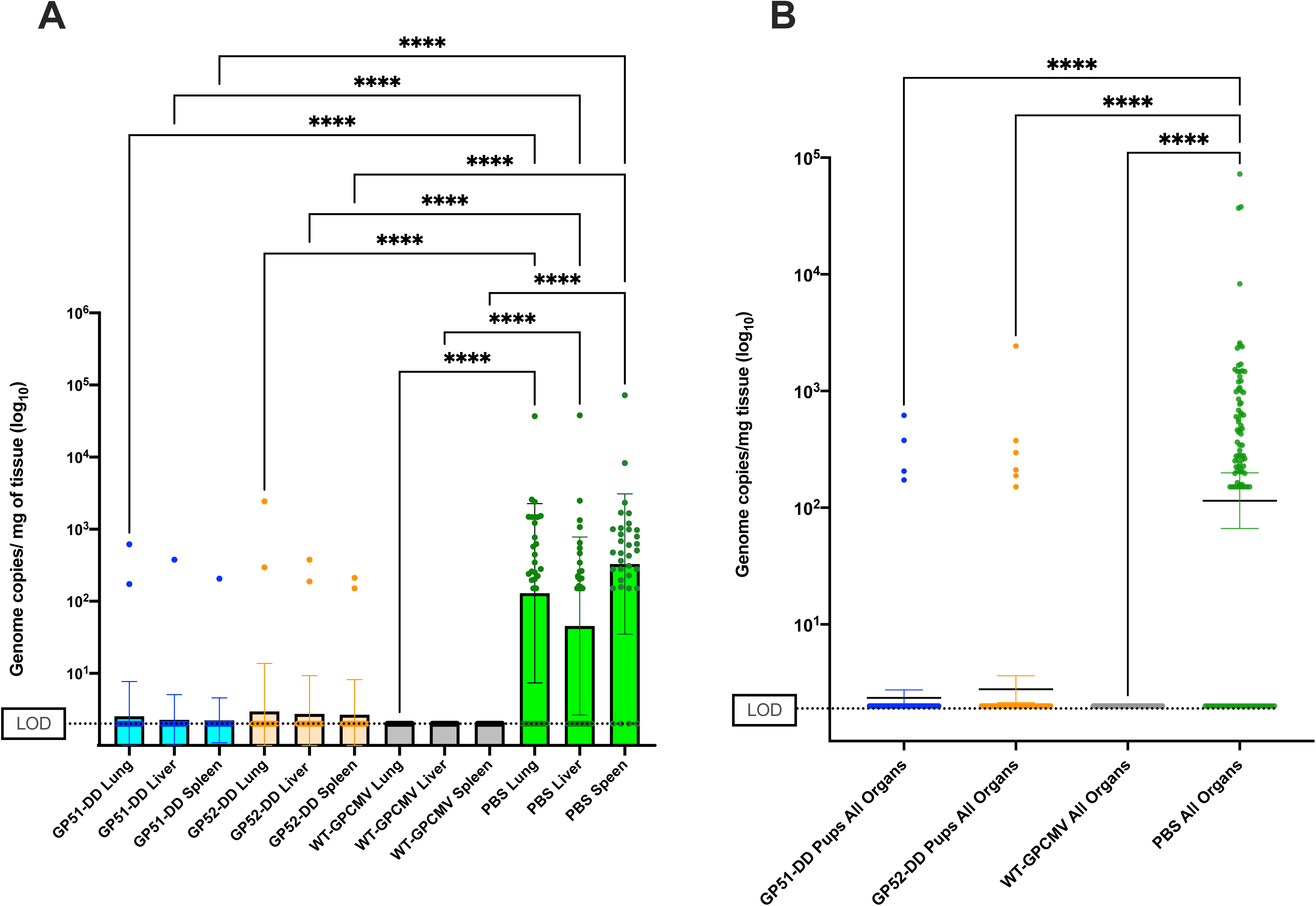
Immunization with GP51-DD or GP52-DD significantly reduces viral loads in pup organs. DNA was extracted from pup blood and tissue after delivery and viral loads were quantified by qPCR. (A) PCR results for pup visceral tissue (lung, liver, spleen), for each vaccine group, are compared by organ. Samples below the LOD for the PCR assay (dashed line) were assigned a value of 2 genome copies/mg for statistical comparisons. (B) Results for all visceral organ samples (lung, liver, and spleen) combined for comparisons of each individual vaccine group. Data shown are means of log_10_ transformed viral load data ± one SD, compared by one-way ANOVA. **** p<0.0001.

## 4. Discussion

A vaccine for the prevention of congenital HCMV infection is a major public health priority (36), but the ideal platform for such a vaccine is uncertain. Subunit vaccines based on the immunodominant gB have been evaluated in multiple clinical trials; these trials have demonstrated no better than ∼45-50% efficacy in preventing primary infection in women of childbearing age (37, 38). Although the correlates of immunity required for protection of the fetus following maternal infection and/or immunization are unknown (39), it may be that a broader repertoire of immune responses – beyond anti-gB responses – are required. Such a goal could be achieved with a whole virus HCMV vaccine. The first HCMV vaccine candidates were comprised of live strain Towne or AD169; both produced by extensive serial passage on fibroblasts (40–43). Although these vaccines were highly attenuated when administered in human clinical trials, a phase 2 trial of the Towne vaccine in young women with children attending group day-care centers – a high risk population for acquisition of a primary HCMV infection – did not demonstrate vaccine efficacy (44). Moreover, a low-passage strain, Toledo, was pathogenic in naturally infected or Towne-vaccinated humans even following administration of as few as 100 pfu (45). Thus, lack of evidence for efficacy and concerns about safety have dampened enthusiasm for further development of live-attenuated HCMV vaccines.

Replication-defective viruses are a vaccine strategy that, in principle, retains the benefits of a live vaccine while ensuring safety. For example, a whole virus GPCMV vaccine rendered replication-defective by a mutation disrupting *GP85,* an essential capsid-encoding gene and the homolog of *UL85,* elicited a broad repertoire of humoral and cellular responses and provided a high level of protection against congenital transmission (46, 47). However, in the rhesus macaque model a whole virus vaccine rendered replication-defective by the deletion of sequences encoding glycoprotein L and several genes involved in immune evasion was less effective than a soluble recombinant gB protein vaccine at reducing post-challenge DNAemia (48). While the replication-defective vaccine was superior in inducing cellular immunity, it elicited less robust humoral immunity, both with respect to viral antigen-reactive antibodies and neutralizing antibodies (48). Thus, at least in the rhesus model, the replication-defective strategy did not confer any clear advantage over the gB subunit vaccine.

A replication-defective HCMV vaccine, V160, was recently evaluated in human clinical trials (21, 49–51). The vaccine is based on the AD169 strain, modified to express UL51, IE1, and IE2 as N-terminal DD fusions. In addition, a mutation in *UL131A* was repaired to restore expression of the PC (21, 50, 51). Safety and immunogenicity of V160 were confirmed in a phase 1 study, and neutralizing antibody titers induced by V160 vaccination in CMV-seronegative individuals were similar to those observed with natural infection (52). This included antibody responses that specifically neutralize epithelial cell entry (52), which were deficient in subjects vaccinated with gB/MF59 subunit vaccine or the live Towne vaccine, presumably because neither are able to induce antibodies specific to the PC (53).

In a recent phase 2b study, a three-dose series of the V160 vaccine demonstrated an efficacy of 44.6% against acquisition of primary HCMV infection in young seronegative women (19). However, because serological detection of naturally acquired infections in the context of antibodies induced by V160 vaccination was not feasible, PCR was used, and surprisingly, a significant number of placebo recipients were PCR-positive but never developed CMV-reactive antibodies. Thus, some PCR-positive V160 recipients may similarly have been false-positives (54). Based on this assumption and correcting for false-positives among the V160 recipients, efficacy increased to 67% or 77%, depending on vaccine group (19).

In this study, we examined two recombinant GPCMV vaccines where essential viral proteins were modified using the same protein destabilization technology as was used in V160. For both the GP51-DD and GP52-DD vaccines, a few animals exhibited low-level DNAemia following immunization. This was not surprising in the case of GP51-DD, which replicates *in vitro* without Shield-1 (Fig. 1B). GP52-DD is much more dependent on the presence of Shield-1 for replication, though very low levels of replication may occur *in vivo* and *in vitro*. In the experiment shown in Fig. 1B, small foci of infected cells were observed in the cell monolayer, and in a replicate experiment at higher MOI very low levels of cell-free infectious virus (∼20 pfu/ml) were detected at 14 days post infection (data not shown). The low-level DNAemia that was detected could also be an artifact of DNA from the vaccine inoculum or DNA produced by cells unproductively infected by virions in the vaccine, that gradually reached the blood. Even so, we consider both GP51-DD and GP52-DD vaccines *replication-deficient* rather than fully replication-defective. As in V160, it may be necessary to target multiple essential viral proteins for degradation in order to generate a fully replication-defective GPCMV vaccine.

Despite evidence that GP52-DD is far more replication-deficient than GP51-DD *in vitro*, GP52-DD was similarly immunogenic to GP51-DD. Both replication-deficient vaccines compared favorably with fully replication-competent WT-GPCMV with respect to the induction of total GPCMV-reactive IgG, IgG avidity maturation, and virus neutralizing activities. Thus, the ability to replicate efficiently *in vivo* is not a requirement for vaccine immunogenicity, at least in so far as the immune parameters that were measured.

Preconception vaccination with WT-GPCMV appeared to induce sterilizing immunity for dams that underwent subsequent viral challenge during pregnancy. While post-challenge DNAemia was also not detected in dams vaccinated with GP51-DD, the presence of viral DNA in two pups born to GP51-DD-immunized dams suggests that GP51-DD-vaccination did not provide complete sterilizing immunity. GP52-DD was partially protective, as 44% of immunized dams exhibited low-level post-challenge DNAemia, while the remaining 66% were negative at all time points. Vaccination with GP52-DD and WT-GPCMV slightly improved pup birthweights compared to sham-vaccination, but no differences were observed with respect to the duration of pregnancy or magnitude of pup mortality, which for all groups were on par with that of unchallenged pregnancies (55).

When pups were examined for evidence of intrauterine infection by PCR, 91% of pups born to sham-inoculated dams were positive in at least one sample, indicating efficient vertical transmission in the absence of preexisting maternal immunity. Vaccination with WT-GPCMV fully prevented vertical transmission, while vaccination with GP52-DD or GP51-DD reduced but did not eliminate transmission, resulting in 9.6% or 5.1% of pups infected, or protective efficacies of 89% or 94%, respectively.

While there are limits to which these findings can inform development of HCMV vaccines, two aspects are worth noting. *First*, immunity induced by WT-GPCMV was more protective against maternal DNAemia than that induced by GP52-DD, and more protective against vertical transmission than either GP51-DD or GP52-DD. However, all three immunizations induced similar humoral responses. This suggests that other aspects of the adaptive response to WT-GPCMV are necessary to prevent 100% of fetal infections. Such components include T-cells (56), as well as non-neutralizing Fc-mediated humoral responses, such as have been proposed for HCMV (57–60). Moreover, as the breadth of humoral responses to the spectrum of GPCMV antigens was not examined, GP51-DD or GP52-DD may have inadequately induced antibodies to specific viral proteins that are important for sterilizing immunity. Thus, further efforts to identify immune parameters that are elevated following WT-GPCMV compared to GP51-DD or GP52-DD vaccination are warranted.

*Second,* while the 89% efficacy against fetal infection by the more replication-deficient GP52-DD vaccine is encouraging, efficacy against maternal infection was low and on par with that reported for V160 in humans (19). As maternal infection will likely be the primary endpoint for approval of a human vaccine (61), there is room for improvement of replication-defective vaccines over the benchmark set by V160. The design of V160 could potentially be improved in several ways. For example, given the plethora of genetic defects inherent in the AD169 background used for V160 (62, 63), one potential improvement could be to base a replication-defective vaccine on a wild type HCMV such as the Merlin strain (64). A second potential deficiency arises from destabilization of IE1 and IE2 through DD fusions (21), as *in vivo*, degradation of IE1 and IE2 likely precludes V160-infected cells from expressing late viral proteins, including many important targets of humoral and cellular immunity. Thus, a replication-defective vaccine in which only late proteins are destabilized might be far more immunogenic.

Lastly, numerous strategies could be employed to further enhance immunogenicity of a replication-defective vaccine, for example, through insertion of gene cassettes to hyper-express certain viral immunogens, or deletion of viral genes that encode proteins having immune-modulatory functions.

Although an HCMV mRNA vaccine is in the advanced stages of clinical trials (3), no vaccine is yet licensed for prevention of HCMV infection in women of childbearing age. Despite sub-optimal efficacy reports from an initial clinical trial (19), efforts to improve immunogenicity and enhance efficacy of replication-defective vaccines are warranted. The results presented here demonstrate that a whole virus GPCMV vaccine that is highly replication-deficient provides significant levels of protection against maternal and congenital infection. Thus, a fully replication-defective GPCMV that emulates V160, while avoiding the IE destabilization that is a feature of that vaccine, could be used as a platform to explore modifications designed to enhance immunogenicity and protective efficacy of vaccine-mediated protection against maternal and fetal infection.

## Acknowledgments

This work was supported by grant 1R01HD098866-01A1 (to M.R.S, A.P.G., and M.A.M) from the National Institutes of Health. M.R.S. acknowledges a Grant-in-Aid Award 740309 from the University of Minnesota Research and Innovation Office. C.E.T. was supported by grant 2T35AI118620-06, a G.E.R.M. grant from the Infectious Diseases Society of America, and a UMN Foundation medical student research grant. A.B.-G. was supported by a grant from the Life Sciences Summer Undergraduate Research Program at the UMN (NHLBI R25HL088728). The authors thank Dai Wang and Merck Vaccines for kindly providing Shield-1.

